# Specific bacterial cell wall components influence the stability of Coxsackievirus B3

**DOI:** 10.1101/2021.05.14.444268

**Authors:** Adeeba H. Dhalech, Tara D. Fuller, Christopher M. Robinson

## Abstract

Enteric viruses infect the mammalian gastrointestinal tract and lead to significant morbidity and mortality worldwide. Data indicate that enteric viruses can utilize intestinal bacteria to promote viral replication and pathogenesis. However, the precise interactions between enteric viruses and bacteria are unknown. Here we examined the interaction between bacteria and Coxsackievirus B3, an enteric virus from the picornavirus family. We found that bacteria enhance the infectivity of Coxsackievirus B3 (CVB3) in vitro. Notably, specific bacteria are required as gram-negative *Salmonella enterica*, but not *Escherichia coli*, enhanced CVB3 infectivity and stability. Investigating the cell wall components of both *S. enterica* and *E. coli* revealed that structures in the O-antigen or core of lipopolysaccharide, a major component of the gram-negative bacterial cell wall, were required for *S. enterica* to enhance CVB3. To determine if these requirements were necessary for similar enteric viruses, we investigated if *S. enterica* and *E. coli* enhanced infectivity of poliovirus, another enteric virus in the picornavirus family. We found that, in contrast to CVB3, these bacteria enhanced the infectivity of poliovirus in vitro. Overall, these data indicate that distinct bacteria enhance CVB3 infectivity and stability, and specific enteric viruses may have differing requirements for their interactions with specific bacterial species.

**Importance:** Previous data indicate that several enteric viruses utilize bacteria to promote intestinal infection and viral stability. Here we show that specific bacteria and bacterial cell wall components are required to enhance infectivity and stability of Coxsackievirus B3 in vitro. These requirements are likely enteric virus-specific as the bacteria for CVB3 differs from poliovirus, a closely related virus. Therefore, these data indicate that specific bacteria and their cell wall components dictate the interaction with various enteric viruses in distinct mechanisms.

## Introduction

Enteric viruses are highly infectious pathogens that account for 1.3 million deaths in neonates and children on an annual basis worldwide (1–3). Enteric viruses initiate infection in the gastrointestinal tract and are spread by contamination in food, water, and direct contact (4, 5). Many enteric viruses belong to the Picornaviridae family, whose members are characterized by a positive sense, single-stranded RNA genome, encapsulated by an icosahedral capsid (6). Coxsackievirus is a common, clinically isolated picornavirus and acts as an etiological agent of Hand, foot, and mouth disease, hemorrhagic conjunctivitis, and viral myocarditis (7–9). Because hygiene acts as an essential barrier and preventative factor against the viral fecal-oral route of transmission, the virus predominantly infects infants, young children and accounts for 11% of the fatality rate in neonates (5, 10, 11). Despite this, there are no vaccines or treatments for Coxsackievirus infections.

The gastrointestinal tract is colonized by more than 10^14^ bacteria (12, 13) which aid in host digestion, regulate gut homeostasis, and protect from pathogenic bacteria (14–16). Imbalances in this microbial niche have been related to a variety of diseases such as diabetes (17), inflammatory bowel disease (IBD) (18), and obesity (19). Previous studies have shown that these intestinal microbes also aid in enhancing replication and pathogenesis of enteric viruses, including Coxsackievirus B3 (CVB3) (20–28) but the mechanism behind these interactions remains unclear.

Previous studies have shown that intestinal bacteria can aid in the stability of multiple enteric viruses (20, 22, 24, 29). Interestingly, the bacteria required to enhance viral stability may differ amongst enteric viruses (21, 30). Here we show that species-specific bacteria improve CVB3 stability, and this stability is dependent on the structure of lipopolysaccharides present on the bacterial cell wall. Furthermore, we confirm that even closely related enteric viruses utilize different bacteria to enhance their stability. Overall, these data indicate that bacterial-mediated enhancement of enteric viral stability may be a broad, conserved feature of bacteria-virus interactions and that species-specific bacteria may regulate this mechanism in different viruses.

## Results

### CVB3 infection is enhanced by intestinal bacteria ex vivo and in vitro

Antibiotic treatment of mice before oral inoculation with CVB3 decreases fecal shedding and lethality compared to untreated mice (31). To assess if fecal bacteria could also enhance CVB3 infectivity, we treated male interferon a/b receptor knockout (*Ifnar^-/-^*) mice with an antibiotic cocktail for ten days to deplete their intestinal bacteria (Fig. 1A). Following antibiotic treatment, fecal samples from antibiotic-treated mice, untreated mice, or phosphate-buffered saline (PBS) were incubated with 10^5^ PFUs of CVB3 at 37°C for 6 hours, and CVB3 infectivity was measured by a standard plaque assay on HeLa cells. We found that CVB3 infectivity was significantly increased when incubated with feces from untreated mice compared to antibiotic-treated or PBS (Fig. 1B). Since previous data suggested that specific bacteria may be required to enhance CVB3 replication and lethality in vivo (21, 30), we next sought to determine if particular bacteria improve CVB3 infectivity in vitro. CVB3 was incubated with selected gram-positive and gram-negative bacteria at 37°C, followed by a viral plaque assay. CVB3 lost viral titer when incubated with DMEM but gained viral titer when incubated with *Bacillus* strains, *Pseudomonas aeruginosa, Enterobacter cloacae*, and *Salmonella enterica* (Fig. 1C). Interestingly, *Lactococcus lactis, Enterococcus faecium, Klebsiella pneumoniae*, and *Escherichia coli* did not enhance the infectivity of CVB3. These data indicate that bacteria enhance CVB3 infectivity in a species-specific manner.

**Figure 1:**
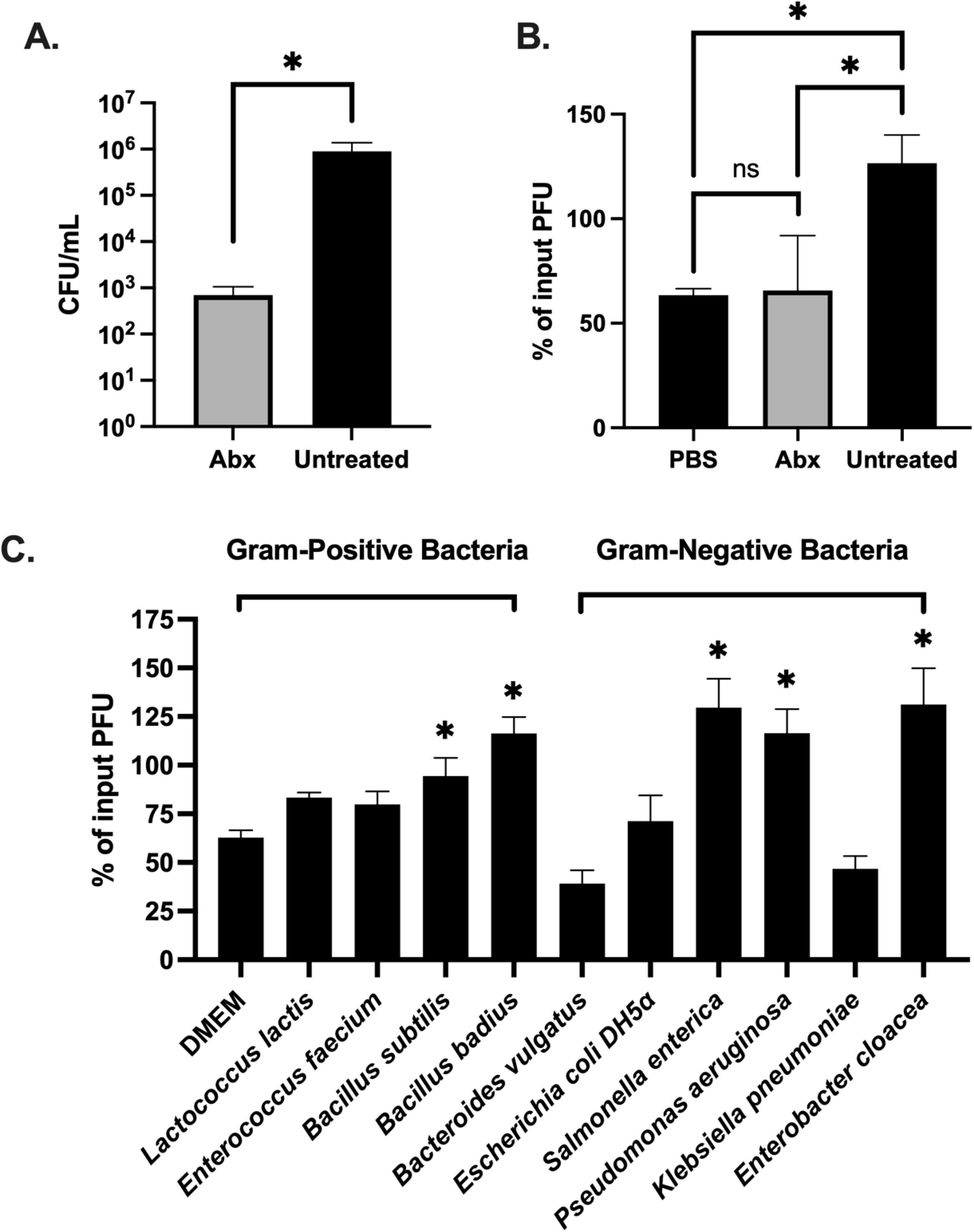
Fecal bacteria enhance the infectivity of CVB3 in vitro. Male C57BL/6 *Pvr^+/+^, Ifnar*^-/-^ mice were treated with an antibiotic cocktail for 10 days. (A) Bacterial loads in the feces. Fecal pellets were collected from the antibiotic-treated and untreated group and plated to determine colony-forming units (CFU) per milligram of feces. Dashed line indicates the limit of detection. (B) CVB3 infectivity after exposure to PBS or feces from uninfected or antibiotic-treated mice (6 hours at 37°C). n= 8 mice per group from 4 independent experiments. (C) CVB3 infectivity after exposure to selected gram-positive and gram-negative bacteria (6 hours at 37°C). Data represent 2-6 independent experiments (n = 4 – 12). For all, bars represent mean +- SEM, *p value<0.05 Student t-test.

### CVB3 can bind to both *S. enterica* and *E. coli*, but only *S. enterica* enhances CVB3 stability

*S. enterica* and *E. coli* are both gram-negative bacteria that are closely related (32). Interestingly, we found that *S. enterica*, but not *E. coli*, could enhance CVB3 infectivity in vitro. Previous data has shown that enteric viruses, including CVB3, could bind to bacteria (30); therefore, we investigated if *S. enterica* or *E. coli* could bind to the CVB3 virion. We incubated 10^5^ PFUs of CVB3 with 10^9^ CFUs of either *S. enterica* or *E. coli* at 37°C for 1 hour to allow for binding. As a control, CVB3 was also incubated in PBS alone or with beads with a similar diameter to bacteria to rule out non-specific binding as previously described (29). Following incubation, the samples were centrifuged at 5,000 x g for 10 minutes to pellet bacteria, washed, and quantified for CVB3 bound bacteria. We found significantly higher levels of CVB3 in the *S. enterica* and *E. coli* pellet compared to PBS and beads (Fig. 2). These data indicate that CVB3 can bind to both *S. enterica* and *E. coli*.

**Figure 2:**
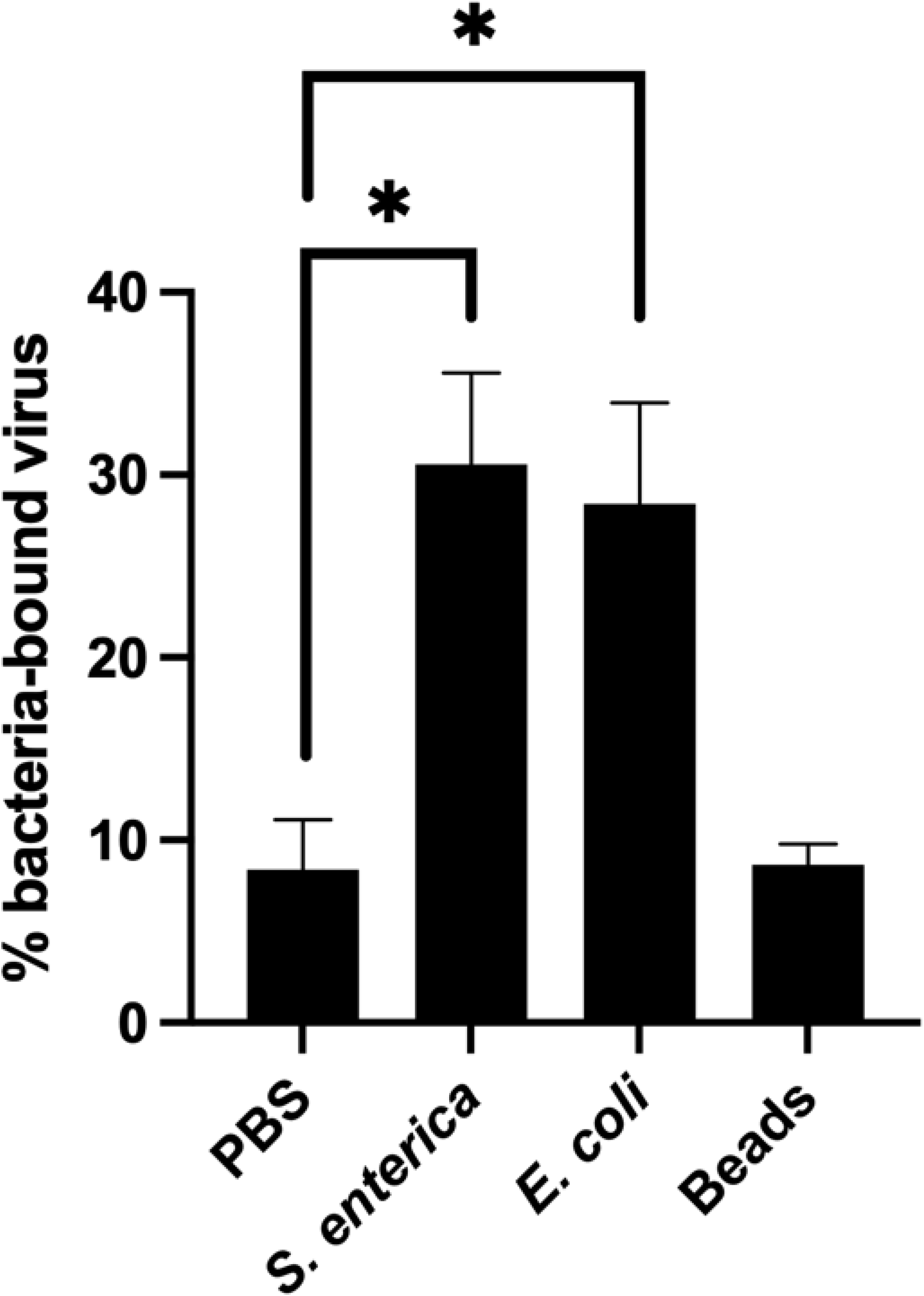
CVB3 bind to *S. enterica* and *E. coli*. 10^5^ PFUs of CVB3 was incubated with 10^9^ CFUs of either *S. enterica* or *E. coli* at 37°C for 1 hour. After incubation, samples were spun down and washed to remove unbound virus. Bound virus was quantified by a plaque assay on HeLa cells. Data represent 3 independent experiments. Bars represent +- SEM, *p value<0.05 Student t-test.

Since CVB3 bound to both *S. enterica* and *E. coli* but only *S. enterica* enhanced infectivity (Fig. 1C), we next quantified the effect of these bacteria on CVB3 stability. To remain infectious, CVB3, like other viruses, must maintain its capsid integrity (6, 33, 34). Previous data indicate that bacteria can improve enteric viruses’ stability, including poliovirus, reovirus, and CVB3 (20, 24, 35); therefore, we sought to determine CVB3 stability by assessing the viral half-life and first-order viral decay in the presence of *S. enterica* and *E. coli* (36). We incubated 10^5^ PFUs of CVB3 with PBS or 10^8^ CFUs of *S. enterica* at 37°C for 0-72 hours. At various times, samples were collected, and CVB3 was quantified by a standard plaque assay. We found that CVB3 incubated with *S. enterica* significantly reduced the decay rate of CVB3 as compared to the virus incubated in PBS (Fig. 3A, B). Further, we found that similar incubation with *E. coli* did not decrease the viral decay rate for CVB3 as compared to PBS, indicating that *E. coli* does not significantly increase viral stability (Fig. 3B). To examine if this effect was dose-dependent, we incubated 10^5^ PFUs of CVB3 at 37°C with PBS or increasing concentrations (10^4^ – 10^8^ CFUs) of bacteria, and viral titers were quantified at various times between 0-72 hours. We observed a significant dose-dependent effect of *S. enterica* on CVB3 stability; however, this stability effect was not observed when CVB3 was incubated with *E. coli* (Fig. 3C and Fig. 3D). These data suggest that *S. enterica*, but not *E. coli*, enhances CVB3 stability in a dose-dependent manner.

**Figure 3:**
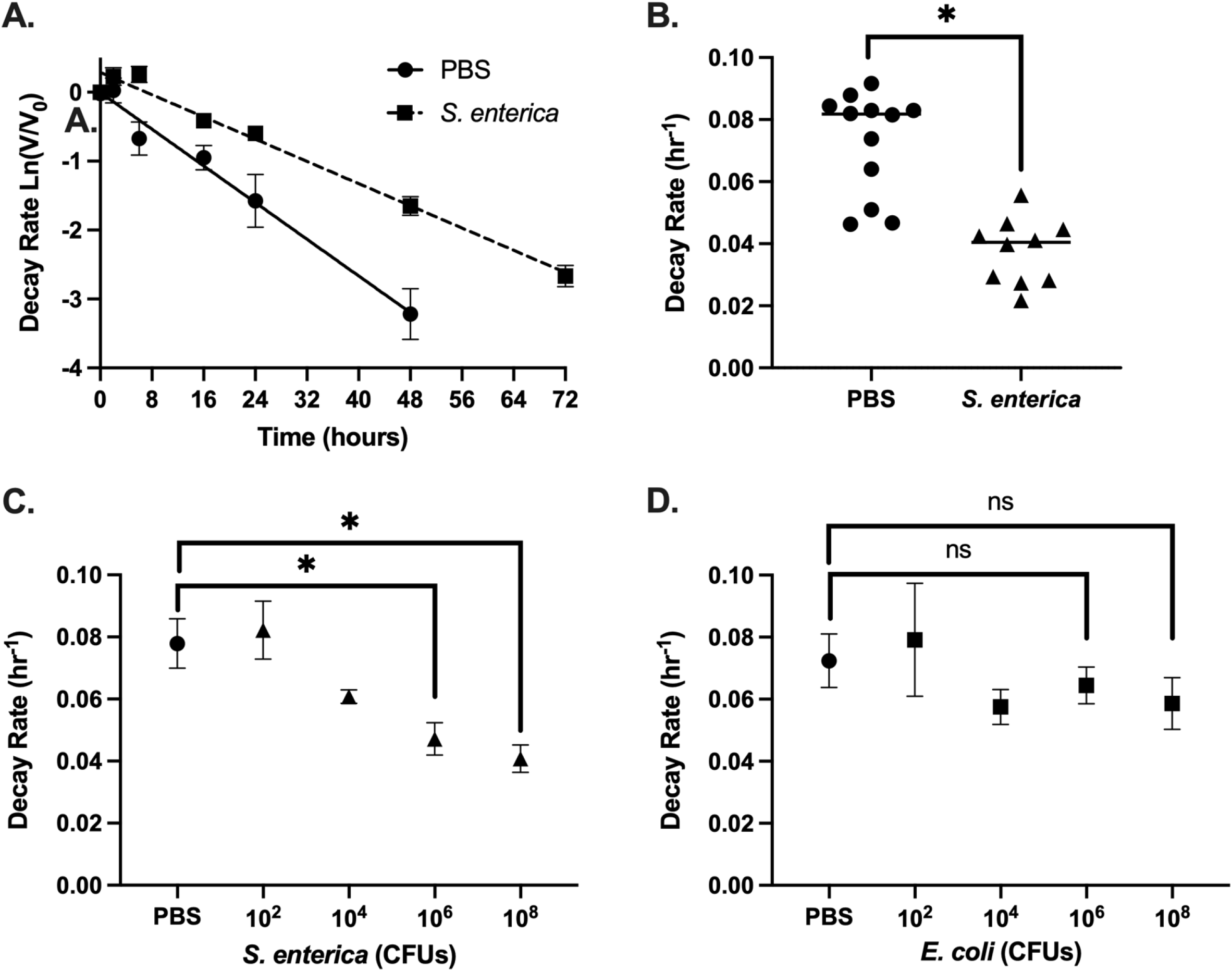
*S. enterica*, but not *E. coli*, enhances CVB3 stability. (A) CVB3 decay at 37°C when incubated with DMEM (slope= −0.06651; R2= 0.6940) or 10^8^ of *S. enterica* (slope= 0.001960; R2= 0.9337). The first-order decay rate for CVB3 was determined by a best-fit line generated by fitted linear regression as previously described (48). (B) Rates of CVB3 Decay in either PBS or 10^8^ CFUs of *S. enterica*. (C) Rates of CVB3 Decay in either PBS or 10^8^, 10^6^, 10^4^, or 10^2^ CFUs of *S. enterica*. (D) Rates of CVB3 Decay in either PBS or 10^8^, 10^6^, 10^4^, or 10^2^ CFUs of *E. coli*. For all, the data represent 4-6 independent experiments, bars represent mean +- SEM *p value<0.05 Student t-test.

### *S. enterica* LPS enhances CVB3 stability, but not *E. coli* LPS

Components of the bacterial cell wall, such as lipopolysaccharide (LPS) and peptidoglycan (PGN), have been shown to bind and stabilize enteric viruses (20, 24). To determine whether bacterial cell wall components could stabilize CVB3, we tested whether LPS from *S. enterica* or *E. coli* could enhance CVB3 stability in vitro. We incubated 10^5^ PFUs of CVB3 with 1mg/mL of LPS from each bacterium or PBS at 42° C for 1 hour and quantified the viral infection on Hela cells. We increased the temperature of our stability assays to 42° C to induce a faster inactivation which has been previously shown as a method to determine the stability of other heat-stable picornaviruses (20). When CVB3 was incubated with either *S. enterica* LPS or *E. coli* K12 LPS, we found the exposure of CVB3 to *S. enterica* LPS, but not *E. coli* LPS, enhanced stability at 42°C (Fig. 4A). We next examined if PGN from *E. coli* could improve CVB3 stability. We found that similar to *E. coli* LPS, PGN from *E. coli* could not enhance CVB3 stability. However, like *S. enterica* LPS, PGN from gram-positive *B. subtilis* could improve CVB3 stability suggesting a bacterial-species specific effect of cell wall components. Finally, we examined if the stability effect of *S. enterica* LPS was dose-dependent. 10^5^ PFUs of CVB3 were incubated with dilutions of *S. enterica* LPS and incubated at 42°C for 1 hour, followed by a plaque assay. We observed a dose-dependent effect of *S. enterica* LPS induced enhancement of CVB3 stability (Fig. 4B). Overall, these data show that bacterial cell wall components LPS and PGN can enhance CVB3 stability in a bacterial species-dependent manner.

**Figure 4:**
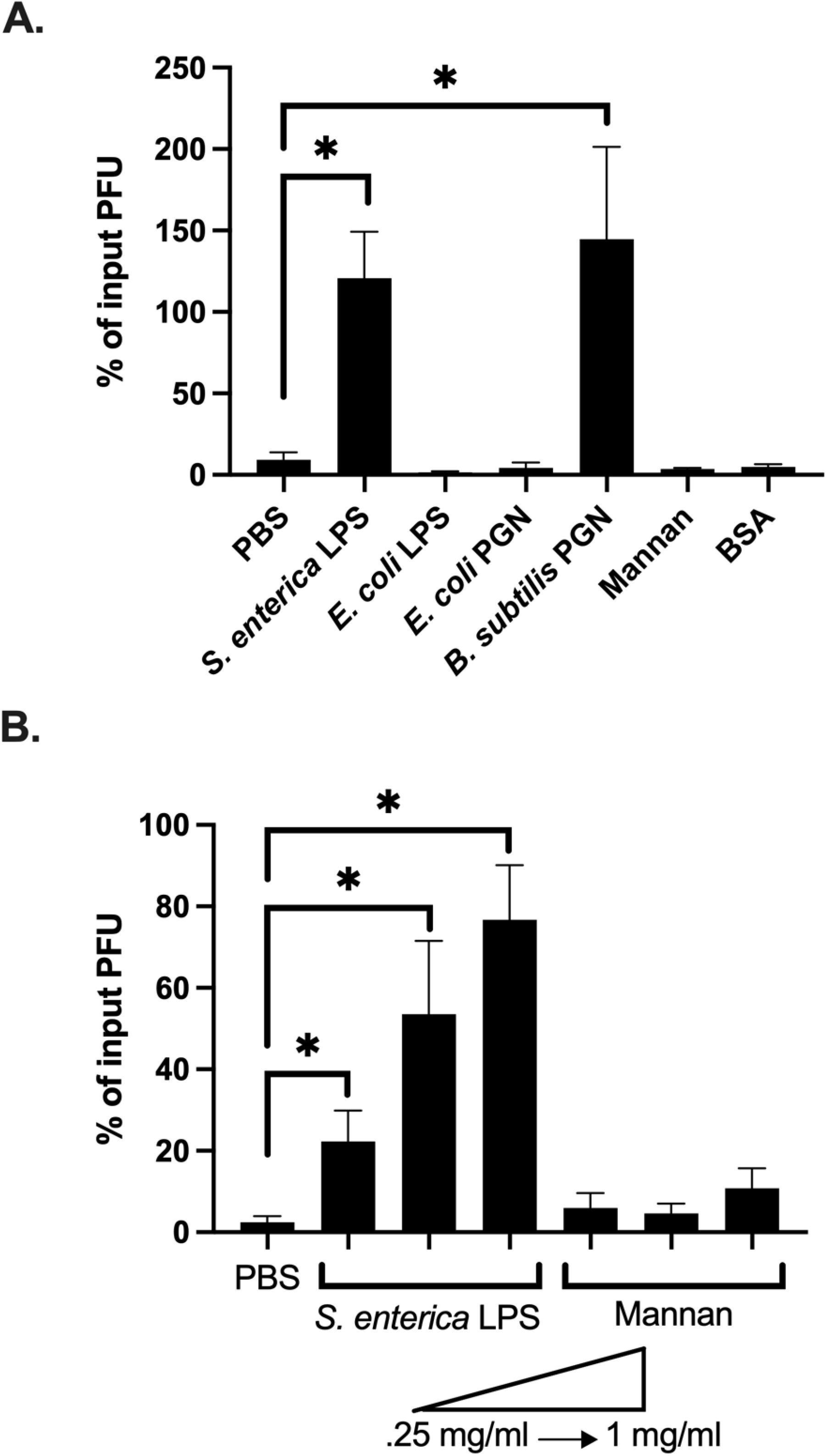
Specific bacterial cell wall components stabilize CVB3. (A) 10^5^ PFUs of CVB3 were incubated in 1 mg/ml concentration of either LPS or PGN from indicated bacterial species at 42°C for 1 hour. PBS, BSA, and Mannan were used as controls. (B) 10^5^ PFUs of CVB3 incubated in increasing concentrations from .25 mg/ml to 1 mg/ml of LPS or mannan PGN at 42°C for 1 hour. For all, data represent 4-5 experiments and bars represent mean +/- SEM. *p value<0.05 Student t-test.

### LPS from *S. enterica* that lacks the outer core and O-antigen does not enhance CVB3 stability

LPS consists of three components: a conserved, hydrophobic lipid A moiety, a semi-conserved hydrophilic core polysaccharide chain, and a polymorphic hydrophilic O-antigen side chain located as the outermost exposed region of LPS (37–40). The O-antigen chain exhibits a strain-specific structural diversity based on the number of repeating oligosaccharide units (41). Smooth gram-negative bacterial strains contain all three components of the LPS structure, while rough bacterial isolates have truncated LPS and lack O-antigen or core components (37, 41) (Fig 5A). Our initial experiments with *E. coli* utilized the rough, lab-adapted DH5a and K12 strains that lack the core and O-antigen components (Fig 3, 4). In contrast, our *S. enterica* strain was a smooth full-length LPS molecule. Therefore, we reasoned that the lack of an O-antigen in *E. coli* might influence CVB3 stability. To test this, we examined the stability of CVB3 in the presence of LPS from *E. coli* 0127, a smooth strain with a fully intact O-antigen. We incubated 10^5^ PFUs of CVB3 with 1mg/mL of LPS from *E. coli* 0127 or PBS at 42° C for 1 hour and quantified the viral infection on Hela cells. Similar to rough strains of *E. coli*, we found that LPS from *E. coli* 0127 did not enhance CVB3 stability (Fig. 5B). Further, CVB3 incubated with whole bacteria from smooth strains of *E. coli* also did not improve infectivity as seen in *S. enterica* (Fig. 5C).

**Figure 5:**
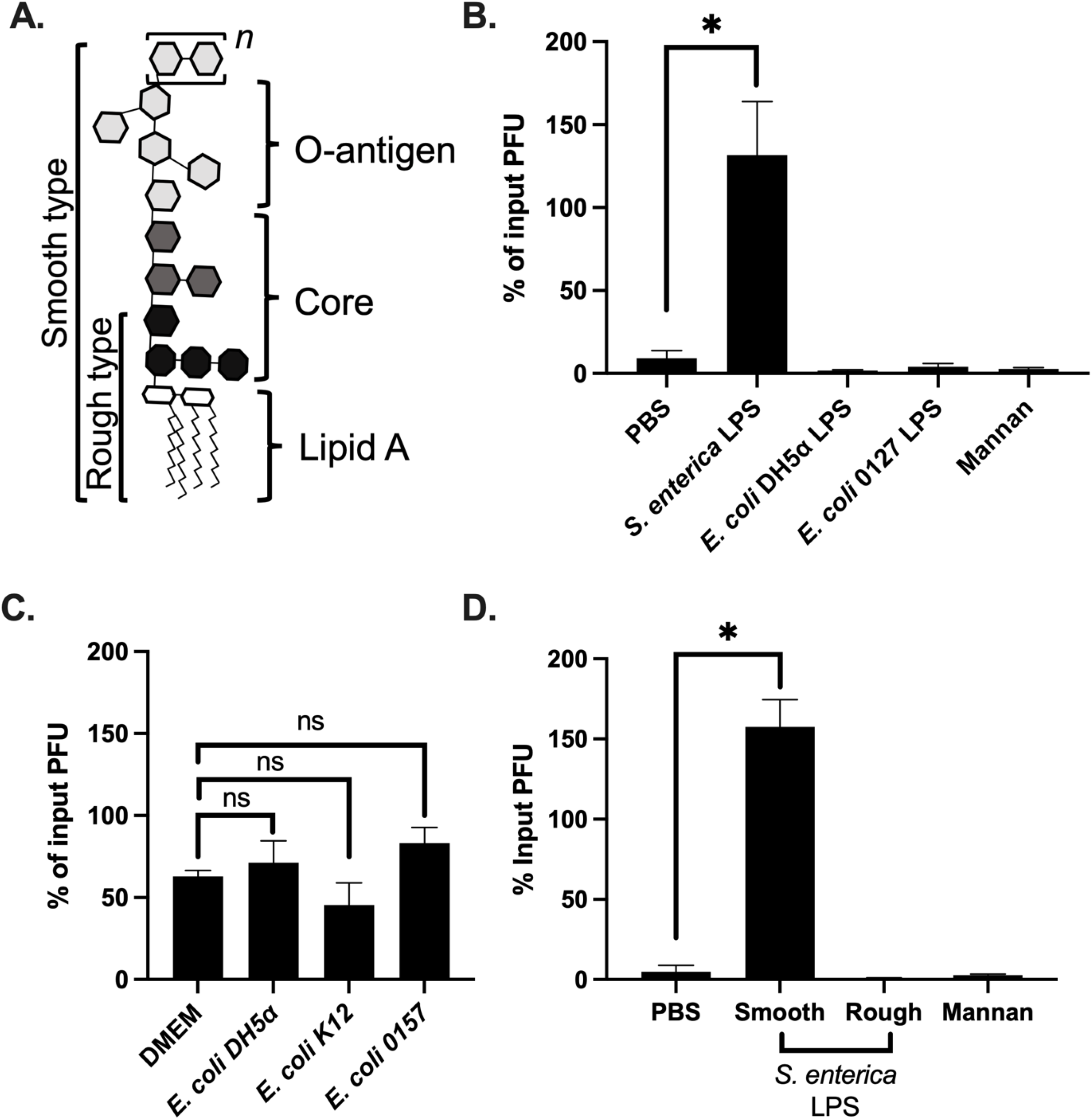
Smooth LPS from *S. enterica* enhance CVB3 stability. (A) The basic structure of bacterial lipopolysaccharide. The O-antigen can have repeating oligosaccharide units (designated *n*). (B) 10^5^ PFUs of CVB3 were incubated in 1mg/ml concentration of LPS from a smooth strain of *S. enterica, E. coli* (0127), a rough strain of *E. coli* (DH5α), or mannan. (C) 10^5^ PFUs of CVB3 incubated with 10^7^ CFUs of rough *E. coli* (DH5α, K12) or smooth *E. coli* (0157) at 37°C for 6 hours. (D) 10^5^ PFUs of CVB3 were incubated with 1mg/mL of LPS from a smooth or a rough strain of *S. enterica Minnesota*. Data represent 4 independent experiments. Bars represent mean +/- SEM *p value<0.05 Student t-test.

Since the O-antigen from *E. coli* did not impact CVB3 stability, we next tested whether the O-antigen or core polysaccharide from *S. enterica* could affect CVB3. 10^5^ PFUs of CVB3 were incubated with 1mg/mL of LPS from smooth or rough strains of *S. enterica Minnesota* at 42°C for 1 hour, and then CVB3 was quantified by a plaque assay. We found that LPS with a fully intact O-antigen from smooth *S. enterica Minnesota* enhanced CVB3 stability, while LPS from the rough strain did not (Fig. 5D). Overall, these results indicate that polysaccharides in the O-antigen or core of *S. enterica* LPS are required to promote CVB3 stability.

### *S. enterica* and *E. coli* enhance the infectivity of poliovirus, a closely related enteric virus

Poliovirus and CVB3 are both members of the Picornaviridae family and share approximately 62.3% sequence identity (21). Previous studies have shown that both poliovirus and CVB3 (20, 22, 29) utilize bacteria to promote infection in vivo; however, the specific bacteria required may differ between the two viruses (21). Further, previous studies indicate that *E. coli* enhances the infectivity of poliovirus (20, 22). Since our data suggest that CVB3 infectivity is enhanced by *S. enterica* and not *E. coli* (Fig 1C), we tested whether *S. enterica* has a stabilizing effect on poliovirus. We incubated 10^5^ PFUs of poliovirus with *S. enterica* or *E.coli* at 37° C for 6 hours, followed by a plaque assay on Hela cells. We found that poliovirus infectivity was significantly enhanced by *S. enterica* and *E. coli* (Fig 6). These data further indicate that highly similar enteric viruses may utilize different bacteria and/or mechanisms to enhance their stability.

**Figure 6:**
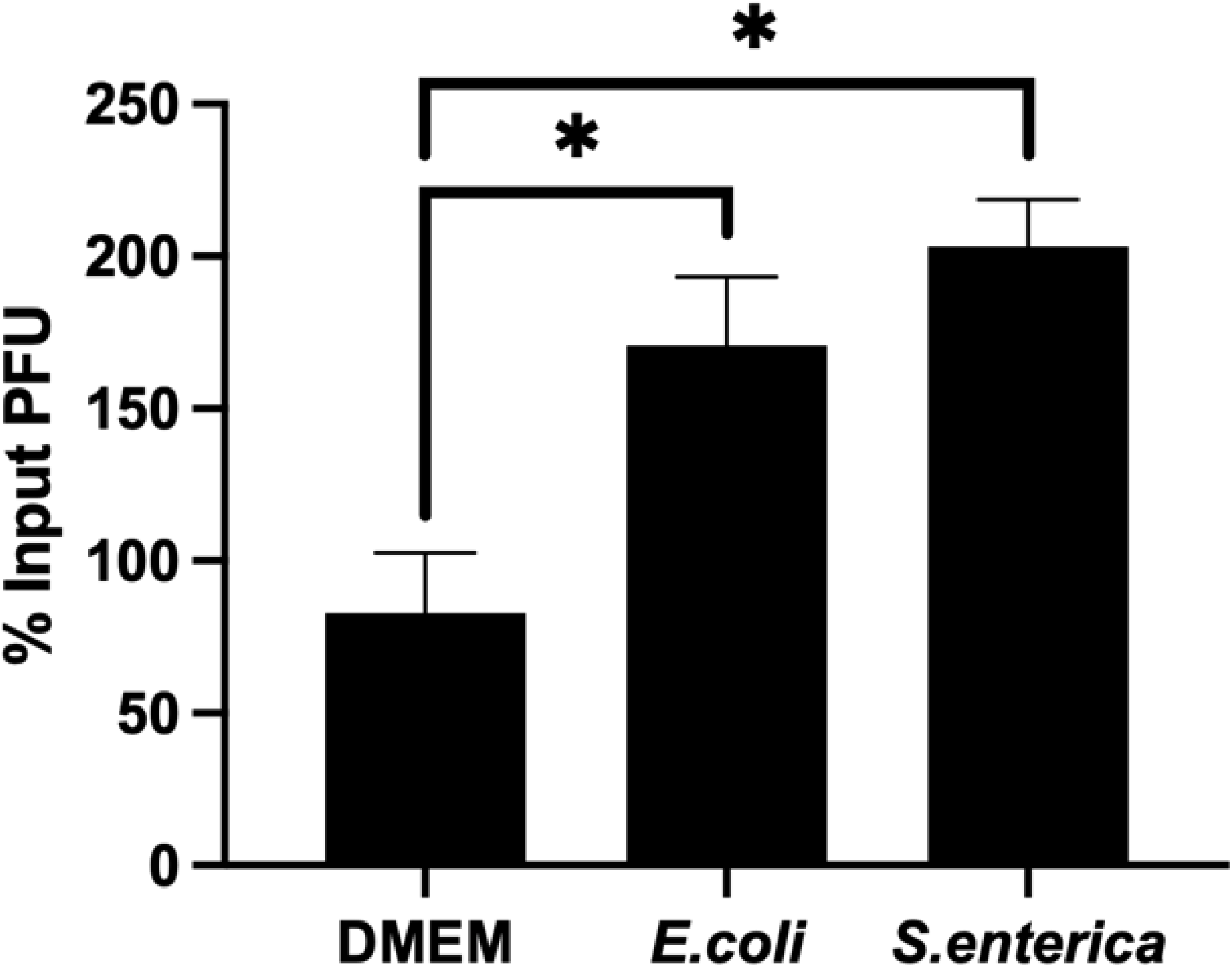
*S. enterica* and *E. coli* enhance poliovirus’s infectivity, a closely related enteric virus. 10^5^ PFUs of PV were incubated with DMEM or 10^7^ of *E. coli* or *S. enterica* at 37°C for 6 hours, followed by a plaque assay on HeLa cells. Data represent 3 independent experiments, bars data represent mean +/- SEM, *p value<0.05 Student t-test.

## Discussion

Coxsackieviruses are spread through the fecal-oral route, where they are exposed to a community of intestinal bacteria. Previous studies have shown the pivotal role intestinal bacteria play in the replication and pathogenesis of several enteric viruses (22–24, 27, 28). Further, data indicate that bacteria can enhance the stability of different enteric viruses, which suggests that this may be a broad, conserved mechanism of viral-bacterial interactions (30, 31). However, the precise mechanisms underlying the different bacterial requirements for interacting with CVB3 remain unclear. Here we show that interactions with specific bacteria and lipopolysaccharide structures enhance CVB3 infectivity and stability.

Our data indicate that gram-positive and gram-negative bacteria can enhance CVB3 infectivity *in vitro* (Fig. 1C). However, we found that closely related gram-negative bacteria, *S. enterica*, and *E. coli*, had different abilities to impact CVB3. In contrast to infectivity, our data indicate that both *S. enterica* and *E. coli* bind to CVB3 (Fig. 2), even though only *S. enterica* could enhance CVB3 stability (Fig. 3). Further, we determined that CVB3 stability was increased when incubated with LPS from *S. enterica*, but not *E. coli* (Fig. 4A and 4B). We hypothesize that the mechanism(s) that facilitate bacterial binding and influence viral stability may differ. In agreement with this hypothesis, data show that poliovirus binding to specific bacteria does not always correlate with an enhancement of infectivity and stability. Of the 41 strains of bacteria that bound to poliovirus, only 30% enhanced poliovirus infection compared to control (29). It is unclear if variances in bacterial cell wall structure account for this observation with poliovirus; however, poliovirus requires at least six subunits of n-acetylglucosamine to stabilize the virion (20, 42). The size of glycan necessary to bind poliovirus differs from the size needed to stabilize, where longer glycans promote virion stability. This length requirement is hypothesized to allow the glycan to bridge two adjacent 5-fold axes to help retain capsid stability (42). Since we found that a smooth *E. coli* strain, which contains n-acetyl glucosamine and is of sufficient length, was unable to stabilize CVB3 (Fig. 5C), these data suggest differences between CVB3 and poliovirus in their interactions with bacteria. In agreement, we found that *S. enterica* and *E. coli* enhanced the infectivity of poliovirus (Fig. 6), but not CVB3. Since previous data revealed that CVB3 and poliovirus have different bacterial requirements to improve replication in vivo (31), and *E. coli* has been shown to enhance reovirus stability (24), these data indicate the bacterial requirements to bind and stabilize enteric viruses may be distinct, and different enteric viruses exploit specific bacterial species to enhance their infection.

*E. coli* and *S. enterica* share several O-serogroups that have identical structures (43–45); however, there are numerous distinct O-antigens present in both species of *S. enterica* and *E. coli* due to lateral gene transfer and side-chain modifications (45). Our data also show that the O-antigen or core polysaccharide from *S. enterica* LPS plays a critical role in stabilizing the CVB3 capsid (Fig. 5B and 5D). In contrast to *S. enterica*, smooth *E. coli* LPS could not stabilize CVB3 (Fig. 4A). This suggests that specific polysaccharides in LPS are required to promote CVB3 stability. Further work is needed to determine whether distinct O-serogroups of *S. enterica* have differing impacts on CVB3. These data also indicate that enteric viral interactions with specific LPS structures may be similar to how bacteriophages bind to LPS as a viral receptor. Bacteriophages bind to unique sugars in LPS which can dictate bacterial tropism (46). We hypothesize that enteric viruses may act similarly; however, it remains to be determined how this impacts infection in the intestine.

In conclusion, our data suggest that binding to specific bacteria through polysaccharides in LPS structures that stabilizes CVB3 virions. We hypothesize that this interaction may help retain infectivity and promote transmission in the environment. Future work will be needed to deduce the specific bacterial LPS sugar(s) required to interact with CVB3. Further, since *S. enterica* is not a common commensal bacterium, the conserved features of the bacterial cell wall will need to be evaluated to identify other bacterial members of the intestinal microbiota that may directly interact with CVB3. Overall, understanding the specific mechanisms for how various enteric viruses utilize bacteria will help develop novel therapeutics and limit future transmission events.

## Materials and Methods

### Cells and virus

HeLa cells, grown in Dulbecco’s modified Eagle’s medium (DMEM) supplemented with 10% calf serum and 1% penicillin-streptomycin at 37°C with 5% CO_2_. CVB3 Nancy infectious clone (IC) was obtained from Marco Vignuzzi (Pasteur Institute, Paris, France) and propagated in HeLa cells as described previously (47). CVB3 was quantified by plaque assays in HeLa cells. Poliovirus (serotype 1, Mahoney) infections and plaque assay were performed using HeLa cells as previously described (20).

### Bacterial strains and cell wall components

*Bacillus* strains, *Pseudomonas aeruginosa, Enterobacter cloacae, Salmonella enterica Lactococcus lactis, Enterococcus faecium, Klebsiella pneumoniae*, and *Escherichia coli* were kindly provided by Ryan Relich (Indiana University School of Medicine). LPS molecules from *E. coli* O127 (L3129), *E. coli* K12 (tlrl-eklps), *Salmonella enterica* (L6011), *Salmonella enterica serotype Minnesota* (L6261, L9764) and Mannan from *Saccharomyces cerevisiae* (M7504) were obtained from Sigma-Aldrich, and PGN molecules from *E. coli* O111 (tlrl-pgneb) and *Bacillus subtilis* (L3265) obtained from Invivogen and Sigma-Aldrich respectively.

### Mouse Experiments

Male mice (C57BL/6 *PVR^+/+^ Ifnar^-/-^*) were administered a combination of 4 antibiotics by oral gavage (ampicillin, neomycin, metronidazole, and vancomycin; 10 mg of each antibiotic per day) for 5 days, followed by the administration in drinking water available *ad libitum* (for ampicillin, neomycin, and metronidazole, 1 g/liter; for vancomycin: 500 mg/liter) for additional 5 days. As a control, a group of mice were left untreated. To confirm bacterial depletion, feces were collected on day 10 of antibiotic treatment. Fecal pellets were weighed and resuspended in 5 times 1xPBS. Fecal bacteria were then quantified by determining CFUs on LB or BHI agar plates.

### Viral infectivity assay

10^5^ PFUs of CVB3 Nancy was incubated with fecal slurry or 1xPBS at 37°C for 6 hours. Following incubation, 1/10^th^ volume of chloroform was added to the samples, centrifuged at 13,000 rcf for 5 mins and supernatant was collected. The viral supernatants were then quantified by a plaque assay on HeLa cells. For incubation with bacteria, specific bacteria were grown in either LB or BHI media in a 5 ml culture incubated at 37°C for 16-20 hours. CFU/ml was quantified by absorbance measurements at OD600. Bacterial cultures were pelleted at 3,250 rpm for 30 mins, resuspended in 1x PBS, and centrifuged again. The pellet was homogenized in 500 uL of DMEM, and 10^7^ CFU of each bacterium, normalized to 200 uL with 1xPBS was added to respective tubes. 10^5^ PFU of CVB3 was added to each bacterial tube and incubated at 37 °C for 6 hours. 10^5^ PFU of CVB3 in PBS, pre, and post-incubation were used as controls. Following incubation, the mixture was chloroform extracted and quantified by a plaque assay on HeLa cells.

### Viral stability and decay rate

*S. enterica* and *E. coli* DH5α were grown in BHI and LB media respectively, for 16 hours at 37 °C and CFUs were quantified by plating on BHI and LB agar, as well as by measuring OD600 absorbance values. Bacteria were centrifuged at 3,250 rpm for 30 minutes to remove media and then washed in 1xPBS and centrifuged again. The pellet was resuspended in 500 uL of 1xPBS. Based on CFU and OD600 values, 1×10^8^, 1×10^6^, 1×10^4,^ and 1×10^2^ CFU of bacteria were added to tubes for each of the following time points and normalized to 200 uL using 1xPBS. 10^5^ PFU of CVB3 was added to each tube and incubated at 37 °C for 0, 2, 6, 16, 24, 48, and 72 hours. Following incubation, 1/10^th^ volume of chloroform was added to the samples, centrifuged at 13,000 rcf for 5 mins and supernatant was collected. The supernatant was used to quantify viral titers by a plaque assay on HeLa cells. For stability experiments, 10^5^ PFU of CVB3 was incubated with 200 uL of either 1 mg/ml, .50 mg/ml or .25 mg/ml of LPS, PGN, 1xPBS, or mannan at 42 °C for 1 hour and then used to quantify viral titers by a plaque assay on HeLa cells.

### Statistical Analysis

Comparisons between control and groups were analyzed using a parried or unpaired two-tailed Student t-test. Kolmogorov-Smirnov test was used to compare CFU/mg between feces from antibiotic and untreated mice. For our stability experiments, a simple linear regression was used to determine the best fit line and the decay rate of CVB3 was determined using the slope of each individual experiment.

## Acknowledgments

We thank Dr. Ryan Relich at Indiana University School of Medicine for providing bacterial isolates for these studies. This work is funded by a K01 DK110216, R03 DK124749, and a Showalter Trust Award from the Indiana Clinical and Translational Sciences Institute to C.M.R.

